# LMI4Boltz: Optimising VRAM utilisation to predict large macromolecular complexes with consumer grade hardware

**DOI:** 10.1101/2025.10.29.684571

**Authors:** Thomas Litfin, Joshua Storm Caley, Katharine A. Michie

**Affiliations:** Structural Biology Facility, Mark Wainwright Analytical Centre, University of New South Wales, Sydney, NSW, Australia; Australian BioCommons, University of Melbourne, Melbourne, Victoria, Australia

## Abstract

AlphaFold2 has revolutionised structural biology by enabling the prediction of protein structures approaching experimental quality. AlphaFold3 extends this framework to support modelling broad biomolecular classes while also reducing the computational cost of prediction. However, AlphaFold3 is distributed with licence conditions which restrict general purpose use. Boltz is a permissive, open-source re-implementation of AlphaFold3, but it is bottlenecked by increased VRAM requirements and requires high-end GPU hardware to model large molecular systems. Here we introduce Low Memory Inference for Boltz (LMI4Boltz) which reduces VRAM requirements using in-place updates, offloading tensors to host memory, careful management of functional scope and aggressive chunking of key operations. Using these strategies, LMI4Boltz increases the token size limit of Boltz-2 by 66.7% without sacrificing prediction accuracy. These optimisations improve the accessibility of Boltz using consumer grade hardware and unlock the ability to model large molecular systems. LMI4Boltz is available at https://github.com/tlitfin/lmi4boltz.

## Introduction

The emergence of AlphaFold2 (Jumper *et al*., 2021) for protein structure prediction has initiated a paradigm shift in computational structural biology. High quality models have been generated for the entire UniProt database (Varadi *et al*., 2022) and dedicated efforts have focused on extending predictions to large scale homo- and hetero-oligomeric complexes (Yu *et al*., 2023). The combinatorial complexity of the potential interactome necessitates an efficient structure prediction model to enable comprehensive screening of candidate interactions.

AlphaFold3 (Abramson *et al*., 2024) has substantially improved the efficiency of prediction workloads while also introducing the ability to model general molecular complexes. AlphaFold3 utilises a lightweight MSA processing module and a downstream diffusion process for generating atomic coordinates to reduce the computational requirements of structure prediction. AlphaFold3 is implemented in Jax and is ‘just in time’ compiled to ensure efficient hardware utilisation.

However, the bespoke restrictions associated with the AlphaFold3 licence has motivated the development of open-source re-implementations with more permissive license conditions (Wohlwend *et al*., 2025; Liu *et al*., 2024; Team *et al*., 2025). Boltz is an AlphaFold3-style model for general purpose molecular structure prediction (Wohlwend *et al*., 2025; Passaro *et al*., 2025). Compared with AphaFold3, Boltz reports competitive accuracy and computational efficiency. However, the Boltz implementation utilises the eager execution of PyTorch and does not benefit from integrated compiler optimisation of memory use. As a result, complex prediction size is limited to ∼1600 tokens using consumer grade hardware (24GB VRAM).

To reconcile the memory disparity, we have introduced several optimisations to improve memory utilisation. For example, the pair representation is updated in-place to avoid temporary update tensors. In addition, several infrequently used tensors are offloaded to host memory to minimise baseline memory overhead. Functional scope is also carefully managed to avoid the temporary duplication of large tensors. Finally, after minimising memory utilisation, chunking is applied to additional model layers to mitigate newly emergent bottlenecks.

## Methods

### Inference settings

The Boltz-2 baseline was run using the 744b4ae commit. max_msa_seqs was set to 4096 for consistency between Boltz-1 and Boltz-2 implementations. All modelling was conducted using expandable segments to allow flexible VRAM utilisation by the PyTorch caching allocator.

### VRAM requirement benchmark

A human ubiquitin sequence (length: 76) was predicted by Boltz with the use_msa_server option. The alignment file was re-used to model an increasing number of ubiquitin subunits until each method registered an out-of-memory error. Experiments were run with a NVIDIA H200 GPU (141 GB VRAM) with per process memory fraction set to 0.17 to simulate 24GB VRAM capacity. Execution time was reported as the best of 3 replicates as recorded by the internal progress bar.

### Prediction accuracy benchmark

Structures from the PDB test set were predicted by each of the proposed methods using the top-1 confidence output from 5 diffusion samples. The pre-generated Boltz-1 input files (Wohlwend *et al*., 2025) were used to maintain consistency with prior work. Prediction accuracy was evaluated by computing the lddt (Mariani *et al*., 2013) and TMscore (Zhang *et al*., 2022) of model outputs against reference structures as implemented in OpenStructure v2.10 (Biasini *et al*., 2013). 8pe3 and 8t4r were excluded due to errors parsing the reference structures.

### In-place operations

Updates to the pair representation are applied in-place to avoid creating a temporary update tensor. This strategy also maintains the output in a compact bfloat16 format. By contrast, Boltz-2 produces some intermediate tensors in full precision due to type promotion and auto-casting of layer norm outputs.

### Tensor offloading

During inference, Boltz instantiates several objects with large memory footprints (e.g. the relative position encoding with size LxLx128, z_init with size LxLx128 and pdistogram with size LxLx64). These objects are used infrequently but persist in GPU VRAM once instantiated. Here we move these objects between host and GPU memory to minimise fixed memory overhead. Similarly, all layers with skip connections maintain parallel copies of the pair representation (LxLx128 size) while computing the update function. Here we push the MSA module skip connection to host memory while computing the required update to reduce peak GPU memory load.

### Function scope management

In Boltz, the function scoping of the trifast implementation creates temporary duplicated copies of the large q, k, v tensors. In practice, this mitigates the theoretical savings of the trifast kernel by increasing VRAM requirements compared with naïve triangle attention. By managing function scope to maintain a single copy of the contiguous q, k, v tensors, the theoretical memory and throughput advantages can be realised. A similar strategy is also used to avoid duplication of the relative position encoding input tensor.

### Aggressive chunking

We added command line parameters to allow interactive adjustment of chunk sizes to resolve bottlenecks as they emerged during benchmarking. In the extreme case, chunk_size_transition_z was set to 32 and chunk_size_tri_attn was set to 64. To minimize memory requirements, we extended the existing MSAFormer pair transition chunking strategy to the PairFormer and pair conditioning layers. Finally, we introduced new chunking functionality to the triangle multiplicative update (triangle_mult_gate_nchunks set to 4).

## Results

Boltz-2 supports predictions with size up to ∼1600 tokens using a GPU with 24 GB VRAM (Fig 1A, B). By offloading key tensors to host memory and eliminating transient full precision operations in the model trunk (+memory), this capability can be increased by almost 50% (Fig 1A, B; >2356 tokens). Despite the overhead associated with moving data between devices, the execution time is also marginally improved when modelling 1596 tokens (Fig 1C). This is achieved by running additional operations in bfloat16 as well as reducing the burden on the caching allocator when approaching the device memory limit. Aggressively chunking operations at key memory bottlenecks (+chunk) further increases the length limit to >2660 tokens (Fig 1A, B) with an associated wall time cost of 8.5% for 1596 tokens (cf Boltz-2). It should also be noted that chunking is only required for large complexes and can be selectively disabled at smaller sizes (<2356 tokens) to maximise overall throughput.

**Figure 1.**
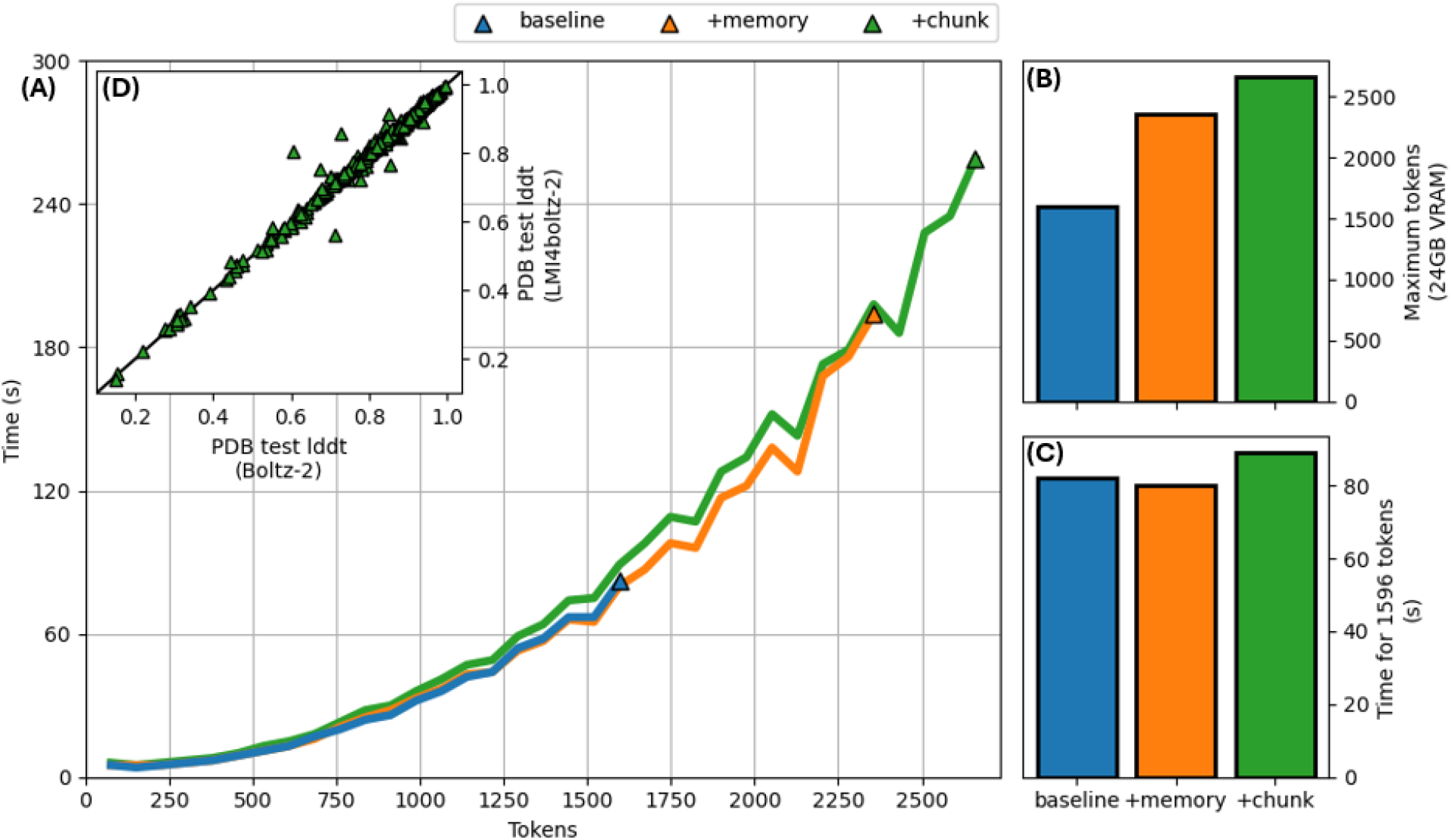
(A) Execution time required to predict the structure of an increasing number of ubiquitin subunits using each of the Boltz-2 implementations. (B) Maximum number of tokens able to be predicted before registering an out-of-memory error with 24GB VRAM. (C) Wall time required to predict the structure of 21 ubiquitin subunits (1596 tokens). (D) lddt of the PDB test set for LMI4Boltz compared with the original Boltz-2 implementation.

LMI4Boltz is particularly efficient for specific sequence lengths which leads to peaks in observed execution times. For example, the time required to predict 2432 tokens is >6% less than the time for a smaller system with 2356 tokens. These optimum lengths coincide with precise multiples of 8 which maximises the efficiency of matrix multiplications on tensor core hardware. This observation highlights a potential strategy to improve throughput for large complexes by padding the length to a multiple of 8 during inference. However, realising this acceleration depends on the nature of the GPU hardware used for prediction.

Despite several modifications, LMI4Boltz outputs closely match those generated by the canonical Boltz-2 implementation. In the PDB test set, the average lddt is 0.820 (cf. 0.820 for Boltz-2). Rare outlier cases occur when the confidence scores of output structures are similar enough that minor fluctuations change the top-1 ranked output (Fig 1D). Using TM-score as an assessment measure highlights outlier complexes which have different binding modes caused by changes in numerical precision (Fig S1A). However, the overall output quality remains unaffected since average TMscore is 0.841 (cf 0.840 for Boltz-2). Exact parity can be restored (Fig S1B) by minor modifications to Boltz-2 including skipping the trivial identity operation in the no dropout setting and casting the outputs of targeted operations to bfloat16 (otherwise promoted to fp32). In addition, since the Boltz-2 implementation of pair transition chunking is not isomorphic to the un-chunked version, the chunk_size_transition_z flag is disabled for LMI4Boltz to restore exact output parity.

## Discussion

In addition to existing optimisations, LMI4Boltz can easily be extended to model larger molecular systems. For example, during triangle multiplicative updates, numerous copies of the pair representation are maintained in memory. These tensors could be juggled between GPU and host as required to relieve a memory bottleneck. However, frequent data movement within an internal loop can significantly increase the overall execution time. Here we have prioritised increasing the sequence length limit while maintaining throughput comparable to the original Boltz implementation. Alternative strategies such as the bespoke chunking strategy introduced in OpenFold (Ahdritz *et al*., 2024) may provide a path to relieve this bottleneck more efficiently.

LMI4Boltz is primarily intended to support inference workloads and modifications such as in-place operations are not compatible with model training. However, the additional reduced precision operations can be directly adapted for training workloads. Offloaded tensors also provide a roadmap for rematerialisation to reduce memory requirements during training (Chen *et al*., 2016). In addition, the memory bottlenecks identified in this work could be optimised in future architectural updates. For example, the pairwise conditioner concatenates the relative position encoding with the trunk pair representation along the channel dimension which spikes memory use. This information can likely be combined more efficiently without impacting model accuracy.

LMI4Boltz significantly increases the sequence length limit of Boltz by careful management of large intermediate tensors and chunking operations at key bottlenecks. While the optimum chunking strategy depends on the available hardware and the size of the prediction system, we demonstrate that robust low-memory inference can be achieved by aggressive chunking for only a small increase in computational cost. Despite minor variations in output compared with Boltz-2 associated with numerical precision, the differences do not negatively impact output quality. Overall, we expect LMI4Boltz to democratise access to the Boltz model by enabling execution using a wide range of consumer-grade hardware.

## Acknowledgements

We gratefully acknowledge the support of the UNSW MWAC Structural Biology Facility and the UNSW Research Technology team for providing access to the computational infrastructure to support this research.

