## Supplementary Material for "LMI4Boltz: Optimising VRAM utilisation to predict large macromolecular complexes with consumer grade hardware"

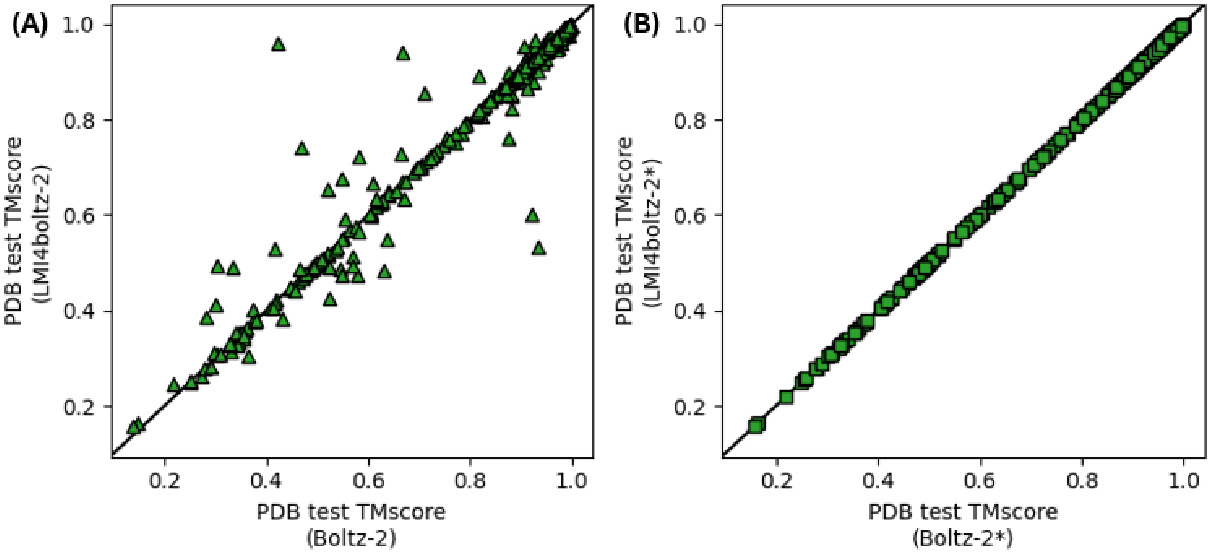


*Figure S1 (A) TMscore of the PDB test set for LMI4Boltz compared with the original Boltz-2 implementation. (B) TMscore of the PDB test set for LMI4Boltz* (without setting chunk_size_transition_z) compared with Boltz-2* (modified to avoid promoting the pair representation to full precision - available in branch 744-parity).*
